# Cyclic Loading Induces Anabolic and Catabolic Gene Expression in ACLs in a Load-Dependent and Sex-Specific Manner

**DOI:** 10.1101/2022.11.11.516153

**Authors:** Lauren Paschall, Sabrina Carrozzi, Erdem Tabdanov, Aman Dhawan, Spencer Szczesny

## Abstract

Anterior cruciate ligament (ACL) injuries are historically thought to be a result of a single acute overload or traumatic event. However, recent studies suggest that ACL failure may be a consequence of fatigue damage. Additionally, the remodeling response of ACLs to fatigue loading is unknown. Therefore, the objective of this study was to investigate the remodeling response of ACLs to cyclic loading. Furthermore, given that women have an increased rate of ACL rupture, we investigated whether this remodeling response is sex specific. ACLs were harvested from male and female New Zealand white rabbits and cyclically loaded in a tensile bioreactor mimicking the full range of physiological loading (2, 4, and 8 MPa). Expression of markers for anabolic and catabolic tissue remodeling, as well as inflammatory cytokines, was quantified using RT-qPCR. We found that the expression of markers for tissue remodeling of the ACL is dependent on the magnitude of loading and is sex specific. Male ACLs activated a tissue remodeling response to cyclic loading below 4 MPa loads but turned off remodeling at 8 MPa. These data support the hypothesis that noncontact ACL injury is a consequence of failed tissue remodeling and inadequate repair of microtrauma resulting from fatigue loading. Conversely, female ACLs downregulate genes responsible for tissue remodeling in response to cyclic loading at all magnitudes, which may explain the increased rate of ACL tears in women. Together, these data provide insight into the remodeling response of ACLs in vivo and potentially offer novel approaches for preventing ACL rupture.

## Introduction

The anterior cruciate ligament (ACL) is one of the most commonly torn ligaments in the human body with tears occurring in more than 200,000 people in the United States each year ^1,2^. The current prevailing scientific and medical perspective is that noncontact ACL tears are caused predominately by a single acute overload or traumatic event ^3^. However, recent studies have provided evidence to suggest that ACLs are susceptible to fatigue failure, where repetitive subfailure loading induces the accumulation of microstructural tissue damage that predisposes the ACL to injury ^4,5^. In addition to direct mechanical damage, data from tendon suggest that fatigue loading of ACLs may also induce tissue degeneration that further weakens the tissue and accelerates fatigue failure ^6^. Previous literature studying tendons demonstrate that fatigue loading and microstructural damage is accompanied by a degenerative cell response characterized by an increase in expression of catabolic proteases (MMPs) and inflammatory cytokines ^7–10^. Given the similarities between tendons and ligaments, it is possible that fatigue-induced degeneration also contributes to ACL rupture. However, to the best of our knowledge, no study has investigated the biological/remodeling response of ACLs to fatigue loading.

Sex-specific ACL remodeling in response to fatigue loading may also help explain the increased susceptibility for ACL ruptures in women. Female athletes are 2-8 times more likely to tear their ACLs compared to their male counterparts ^11,12^. Numerous studies have investigated potential reasons for this sex-specific risk in ACL rupture, which include anatomical, neuromuscular, and hormonal differences between men and women ^13–15^. Previous literature also suggests that there may be sex-specific differences in soft tissue remodeling in response to repetitive loading. For example, estrogen regulates collagen production and crosslinking within ligaments ^13,16–19^ as well as the expression of many MMPs and their inhibitors ^20–23^, which are all responsible for the remodeling of collagenous tissues. Additionally, studies suggest that the overall effect of estrogen on ACL remodeling is negative. Specifically, women are more likely to tear their ACLs during the preovulatory phase (high estrogen level) compared to their postovulatory phase (low estrogen levels) ^24^. Furthermore, estrogen has been shown to reduce ACL strength in rabbits, where ovariectomized rabbits that were treated with estrogen supplements had a lower failure load compared to ovariectomized rabbits without estrogen supplements ^25^. Together, these data suggest that females may have an impaired remodeling response to fatigue loading that could increase the risk of ACL rupture.

Therefore, the objective of this study was to investigate the remodeling response of ACLs to cyclic loading and to see if this response is sex specific. We hypothesized that ACLs will exhibit a dose response to cyclic loading, with lower load magnitudes initiating an anabolic response and higher loads initiating tissue degeneration (i.e., catabolic and inflammatory gene expression). We also hypothesized that female samples will exhibit an increased catabolic response to mechanical loading compared to male samples, which would be consistent with increased ACL rupture rates in women.

## Materials & Methods

### ACL Harvest

Male and female white New Zealand rabbits (2.8-3.2 kg and 14-16 weeks old) were euthanized, and the ACLs were isolated under sterile conditions from an approved IACUC study. The ACLs were kept moistened with phosphate-buffered saline (PBS) during the dissection. The femur and tibia were cut down into bone blocks (approximately 1 x 1 x 1 cm), which were used to grip each sample during mechanical loading. The major and minor diameters and initial length (bone-to-bone) of the ACL were determined using calipers. Sample cross-sectional area was determined by assuming an elliptical cross-section.

### Mechanical Loading

ACLs were placed in a custom-made tensile bioreactor ^26^ with culture media (low-glucose DMEM, 2% PSF, 5% FBS, 25 mM HEPES, 4mM GlutaMAX, 1 mM 2-Phospho-L-ascorbic acid trisodium) and kept at 37 °C and 5% CO_2_. Since underloading the ligament is detrimental to tissue homeostasis ^27,28^, a 0.1 MPa static load was applied for 18 hours, which was sufficient to allow the ACL explants to acclimate to the culture conditions (**Fig. S-1**). After this acclimation period, the bioreactor cyclically loaded the samples to 2 MPa, 4 MPa, or 8 MPa at 0.5 Hz for 8 h. For control samples, a 0.1 MPa static load was maintained for the same duration. These stress levels were determined by scaling prior measurements of in vivo ACL forces within goats during different levels of activity (e.g., standing, walking, and trotting) (Holden et al. 1994).

### Macroscale Tissue Strain

Macroscale tissue strain was calculated as previously described in ^26^. Briefly, tissue displacement was during loading using a linear variable differential transformer (LVDT), and the tissue strain was calculated by dividing the displacement by the initial length of each ACL at the end of the 8-hour loading period.

### Cell Viability

After loading at each condition, ACLs (n = 5-7) were stained with 8 μg/mL fluorescein diacetate (FDA) and 5 μg/mL propidium iodide (PI) in PBS for 10 min at room temperature to visualize live and dead cells, respectively. The ACLs were then sharply dissected from the bone blocks.

Volumetric image stacks were acquired (2.49 μm/pixel) at 3-5 locations along the sample length using an inverted confocal microscope (Nikon AR1 HD). The image stacks were segmented into three different regions based on imaging depth into the tissue (surface: 0 μm depth; middle 32-39 μm depth; deep 56-63 μm depth) and then max z-projected. The images were manually thresholded to eliminate background signal. Cell viability was determined by dividing the area of FDA signal by the total (FDA + PI) signal.

### Gene Expression Analysis

After loading, samples (n = 5-9) were immediately removed from the bioreactor, and the ACLs were sharply dissected from the bone blocks. Freshly harvested samples (n = 4-6) were also collected as a control to evaluate the effect of ex vivo culture on gene expression. The ACLs were rinsed with ice cold RNase-free water and flash frozen in liquid nitrogen. The tissue was then pulverized (6775 Freezer/Mill SPEX) and the total RNA was extracted with RNeasy minicolumns (RNeasy Fibrous Tissue Kit, Qiagen) and quantified using a Qubit 4 Fluorometer (Thermo Fisher). cDNA was synthesized from 10 ng of the total RNA with a commercially available kit (High-Capacity cDNA Reverse Transcription Kit, Thermo Fisher). Quantitative PCR (qPCR) was performed using Taqman probes to measure the expression of anabolic (*COL1A1, COL1A2, LOX, COL3A1, TGFβ1 ACTA, TIMP1, TIMP3*), catabolic (*MMP1, MMP2, MMP10, MMP13*) and inflammatory (*IL-1β, PTGS2*) genes with *ACTB* as the reference gene. Gene expression was quantified relative to the sex-specific statically-loaded control samples using the delta-delta Ct method after correcting for primer efficiencies via the following equation ^29^.

Primer Efficiency: 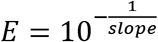 where slope = slope of the Ct vs log(concentration) plot of the standard curve.

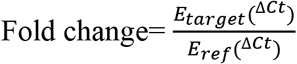

To determine baseline sex-specific gene expression differences, the female freshly harvested and static control samples were compared to the respective male control samples.

### Statistical Analysis

Kruskal-Wallis tests were used to determine the difference in cell viability between loading conditions. Friedman tests were used to determine the difference in cell viability across imaging depth for each condition. Single sample Wilcoxon tests were used to determine differential expression of each gene compared to the respective control condition. To determine if there was an effect of the loading magnitude or sex on gene expression, the expression data for each gene was normalized using a log transformation, and a two-way ANOVA was conducted. Post-hoc Mann Whitney tests were conducted on the non-normalized data to compare the gene expression between loading levels within the same sex and to compare the gene expression between sex at the same load. Mann-Whitney tests were used to determine the difference in cross-sectional area and length between male and female ACLs. Mann Whitney tests were used to determine if there was an effect of sex on macroscale tissue strains at each loading level. Statistical significance was set a p < 0.05. All statistics were performed using the GraphPad Prism (9.3.1) software.

## Results

### Loading protocol maintains cell viability

Some cell death was observed in the explants, with the most death occurring at the tissue surface for each condition and with cell viability increasing with increasing tissue depth (**Fig. S-2 A-D**). In the deep portion of the tissue, there was over 80% viability for the static load and the 2 and 4 MPa cyclically loaded samples (**Fig. 1**). However, cell viability was only 63% for the ACLs cyclically loaded to 8 MPa. While there was no statistically significant differences in cell viability across loading conditions, the data indicated a trend toward lower viability at higher loads (p = 0.07). We also compared cell viability between male and female rabbit ACLs cyclically loaded to 4 and 8 MPa. Female ACLs had a cell viability of 81% and 73% in the deep portion of the tissue at 4 and 8 MPa, respectively (**Fig. S-2E**), which was comparable to the viability of the male ACLs.

**Figure 1:**
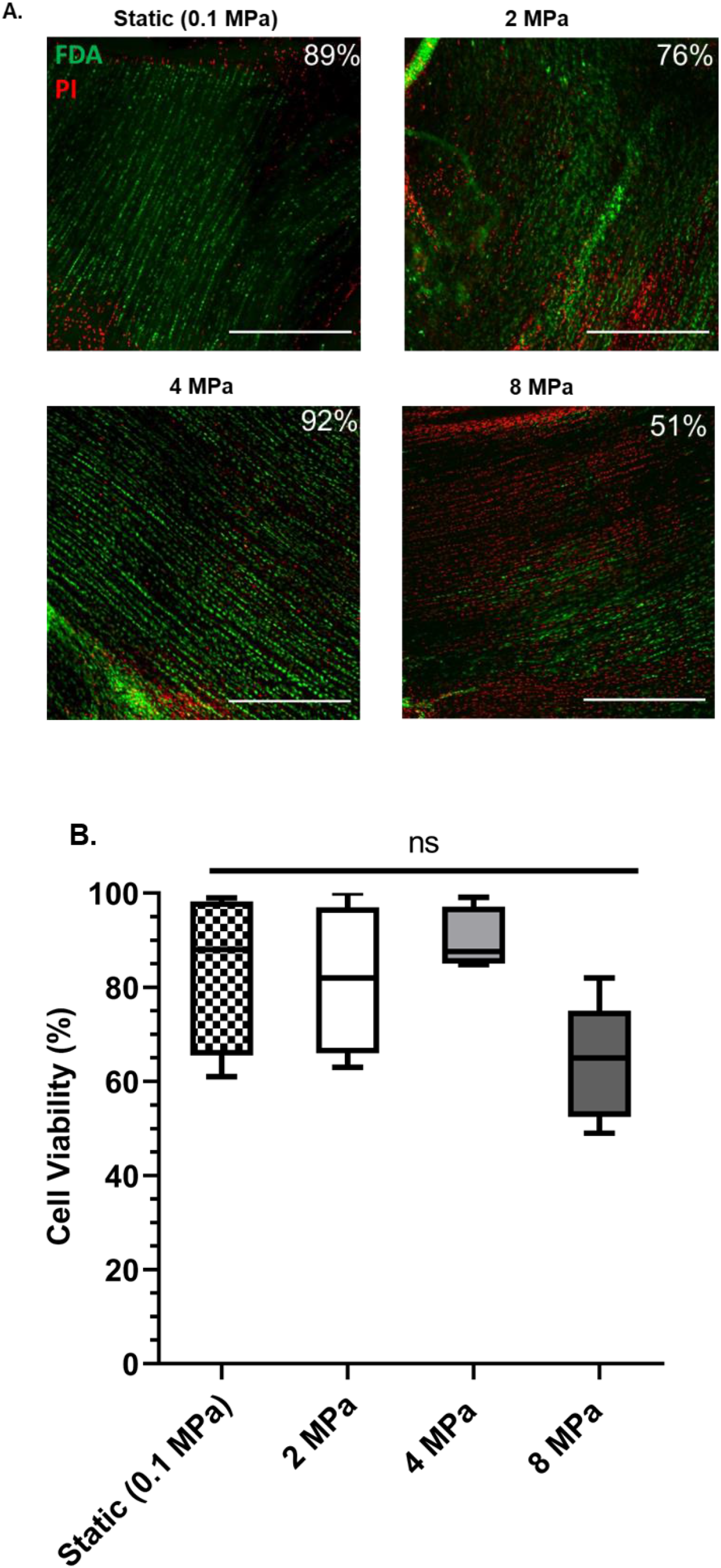
Loading protocol maintains cell viability. (A) Representative images in the deep portion of the tissue for each loading condition. Green (FDA) represents the live cells, red (PI) represents the dead cells (scale bar = 500 um). (B) Quantification of viability for each condition in the deep portion of the tissue (n = 6 for static, n = 7 for 2 MPa, n = 5 for 4 MPa, and n = 5 for 8 MPa). No significant difference in viability across any condition (p = 0.07), evaluated by Kruskal-Wallis test. Data represented as box and whisker plots with the whiskers representing the 10^th^ and 90^th^ percentile of data.

### ACLs exhibit sex-specific gene expression at baseline and in response to mechanical loading

In the freshly harvested ACLs, multiple anabolic (*COL1A1, COL1A2, COL3A1, TGFB1, TIMP1, TIMP2*), catabolic (*MMP1*, *MMP2*), and inflammatory (*PTGS2*) genes were significantly upregulated in the female ACLs relative to the male ACLs (**Fig. 2**). Similarly in the statically loaded ACLs, multiple anabolic (*COL1A1, COL1A2, COL3A1, TIMP1, TIMP2*), catabolic (*MMP1, MMP2*), and inflammatory markers (*IL1β, PTGS2*) were significantly upregulated in the female ACLs relative to the male ACLs. Notably, most genes exhibited similar expression levels between the freshly harvested and statically loaded samples with *ACTA2* and *MMP1* as the only exceptions.

**Figure 1:**
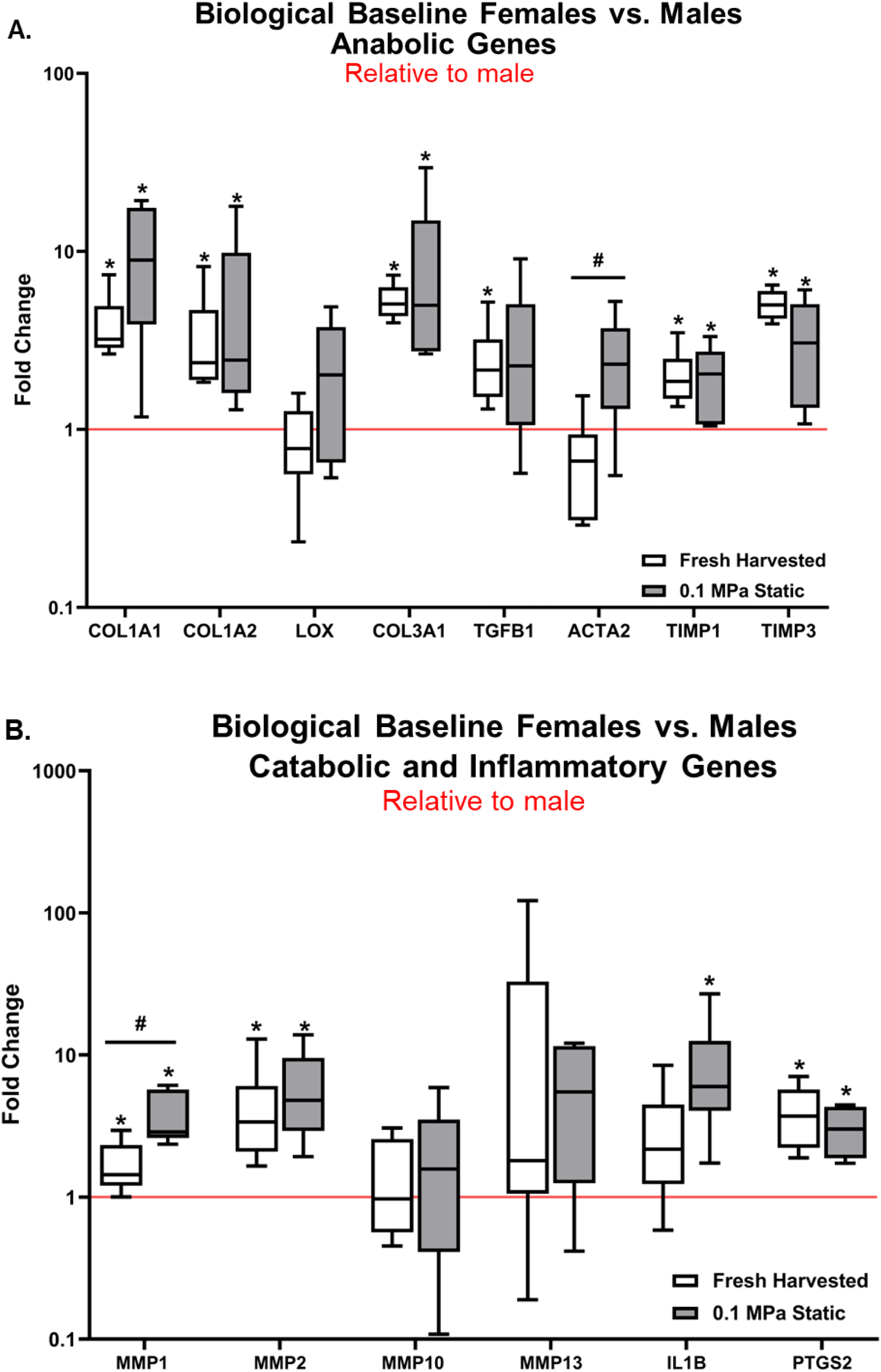
Female ACLs have greater remodeling potential at baseline. RT-qPCR analysis of female ACLs freshly harvested and under 0.1 MPa static load relative to respective male samples (represented by the red line at 1). (A) Anabolic markers. (B) Catabolic and inflammatory markers. (n = 4-6 for fresh harvested and n = 6 for 0.1 MPa static) * p < 0.05 evaluated by one sample Wilcoxon test. # p < 0.05 evaluated by Mann Whitney test. Data represented as box and whiskers plot with the whiskers representing the 10^th^ and 90^th^ percentile of data.

A two-way ANOVA was conducted on the log transformed gene expression data and found that there was a statistically significant effect of load and sex on the expression of nearly all the genes that were investigated (**Table 1**). The only genes in which the effect of loading was not sex dependent were *LOX, ACTA2, PTGS2;* however, there was a significant interaction term for *LOX,* which exhibited sex-specific effects at 2 and 4 MPa but not 8 MPa (**Fig. S-3**).When male ACLs were cyclically loaded to 2 MPa, there was significant upregulation of multiple anabolic (*COL1A2, TGFβ1, TIMP3*), catabolic (*MMP1*), and inflammatory (*IL1β*) markers (**Fig. 3**). Similarly, when male ACLs were cyclically loaded to 4 MPa, there was significant upregulation of multiple anabolic (*COL1A1, LOX, COL3A1, TGFβ1, TIMP1, TIMP3*), catabolic (*MMP2*), and inflammatory (*IL1β*) markers. Only *MMP2* expression was significantly increased between 4 and 2 MPa. When male ACLs were cyclically loaded to 8 MPa, there was significant downregulation of multiple anabolic (*LOX*, *TGFβ1, ACTA2, TIMP1, TIMP3*), catabolic (*MMP1, MMP10*), and inflammatory markers (*PTGS2*) compared to static controls. Only *COL3A1* and *IL1β* was still significantly upregulated at 8 MPa. Comparing the data between 4 and 8 MPa, we observed significantly lower expression for multiple anabolic (*COL1A1, LOX, COL3A1*, *TGFβ1, ACTA2, TIMP1, TIMP3*), catabolic (*MMP1, MMP2*), and inflammatory (*IL1β, PTGS2*) markers.

**Table 1:** P-values resulting from a two-way-ANOVA conducted across all loading conditions and sexes.

| Gene | Load Magnitude Effect | Sex Effect | Interaction |
| --- | --- | --- | --- |
| COL1A1 | 0.011 | <0.0001 | 0.5712 |
| COL1A2 | 0.0029 | <0.0001 | 0.2525 |
| LOX | 0.0008 | 0.92192 | 0.005 |
| COL3A1 | <0.0001 | <0.0001 | 0.8091 |
| TGFB1 | <0.0001 | 0.0007 | 0.2982 |
| ACTA2 | 0.00006 | 0.2542 | 0.7757 |
| TIMP1 | 0.0036 | 0.0043 | 0.064 |
| TIMP3 | <0.0001 | <0.0001 | 0.0286 |
| MMP1 | <0.0001 | 0.0022 | 0.2567 |
| MMP2 | <0.0001 | <0.0001 | 0.0085 |
| MMP10 | <0.0001 | 0.0185 | 0.8377 |
| MMP13 | 0.4274 | 0.0008 | 0.4686 |
| IL1B | 0.034 | <0.0001 | 0.1546 |
| PTGS2 | <0.0001 | 0.2067 | 0.2703 |

**Figure 3:**
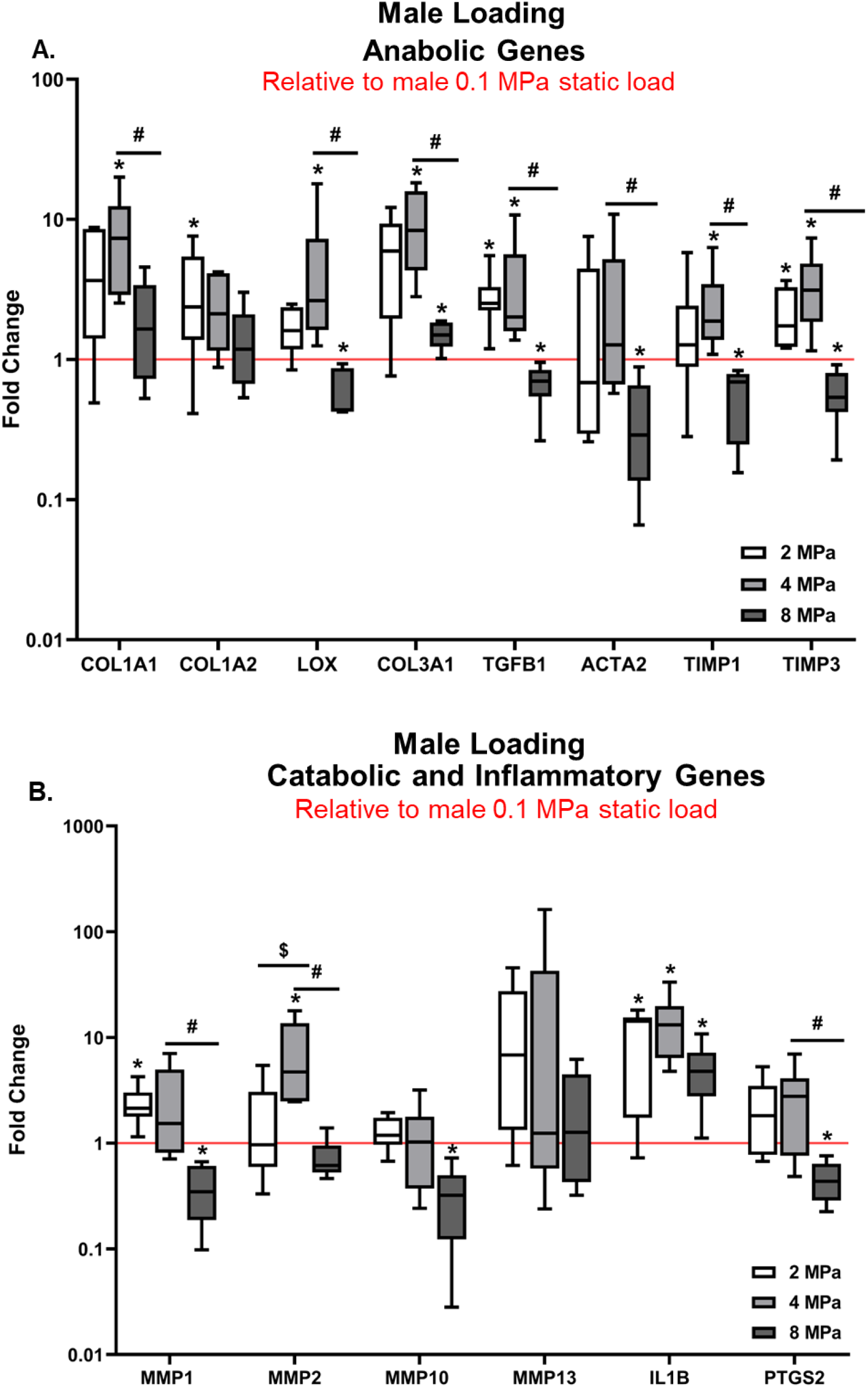
Male ACLs are mechanosensitive in a dose dependent manner. RT-qPCR analysis of male ACLs after being cyclically loaded to 2, 4, and 8 MPa relative to the 0.1 MPa static load (represented by the red line at 1). (A) Anabolic markers. (B) Catabolic and inflammatory markers. (n = 6-9 for 2 MPa, n = 5-6 for 4 MPa, and n = 6 for 8 MPa) * p < 0.05 evaluated by one sample Wilcoxon test). $ represents statistical difference between 2 MPa and 4 MPa (p < 0.05 evaluated by unpaired Mann Whitney test). # represents statistical difference between 4 MPa and 8 MPa (p < 0.05 evaluated by Mann Whitney test). Data represented as box and whiskers plot with the whiskers representing the 10^th^ and 90^th^ percentile of data.

Interestingly, a different response to mechanical loading was seen in the female ACLs (**Fig. 4**). When female ACLs were cyclically loaded to 2 MPa, there was significant downregulation of multiple anabolic (*COL1A1, COL1A2*) and catabolic (*MMP2*) markers. Only *LOX* and *MMP10* were significantly upregulated compared to static controls. When female ACLs were cyclically loaded to 4 MPa, there was also significant downregulation of multiple anabolic (*COL1A1, COL3A1, TIMP1*) and catabolic (*MMP1, MMP2*) markers compared to static controls. Furthermore, anabolic (*LOX*, *TIMP3*) and catabolic (*MMP1*) markers were reduced at 4 MPa compared to 2 MPa. When female ACLs were cyclically loaded to 8 MPa, there was significant downregulation of multiple anabolic (*COL1A1, COL1A2, LOX, COL3A1, TGFβ1, ACTA2, TIMP1, TIMP3*) catabolic (*MMP1, MMP2, MMP13*), and inflammatory (*PTGS2*) markers compared to static controls. Consistent with pervasive downregulation of gene expression at 8 MPa, several anabolic (*COL1A1, COL1A2, LOX, COL3A1, TGFβ1, ACTA2*), catabolic (*MMP2, MMP10*), and inflammatory (*IL1β*) markers had significantly lower expression when cyclically loaded to 8 MPa compared to 4 MPa.

**Figure 4:**
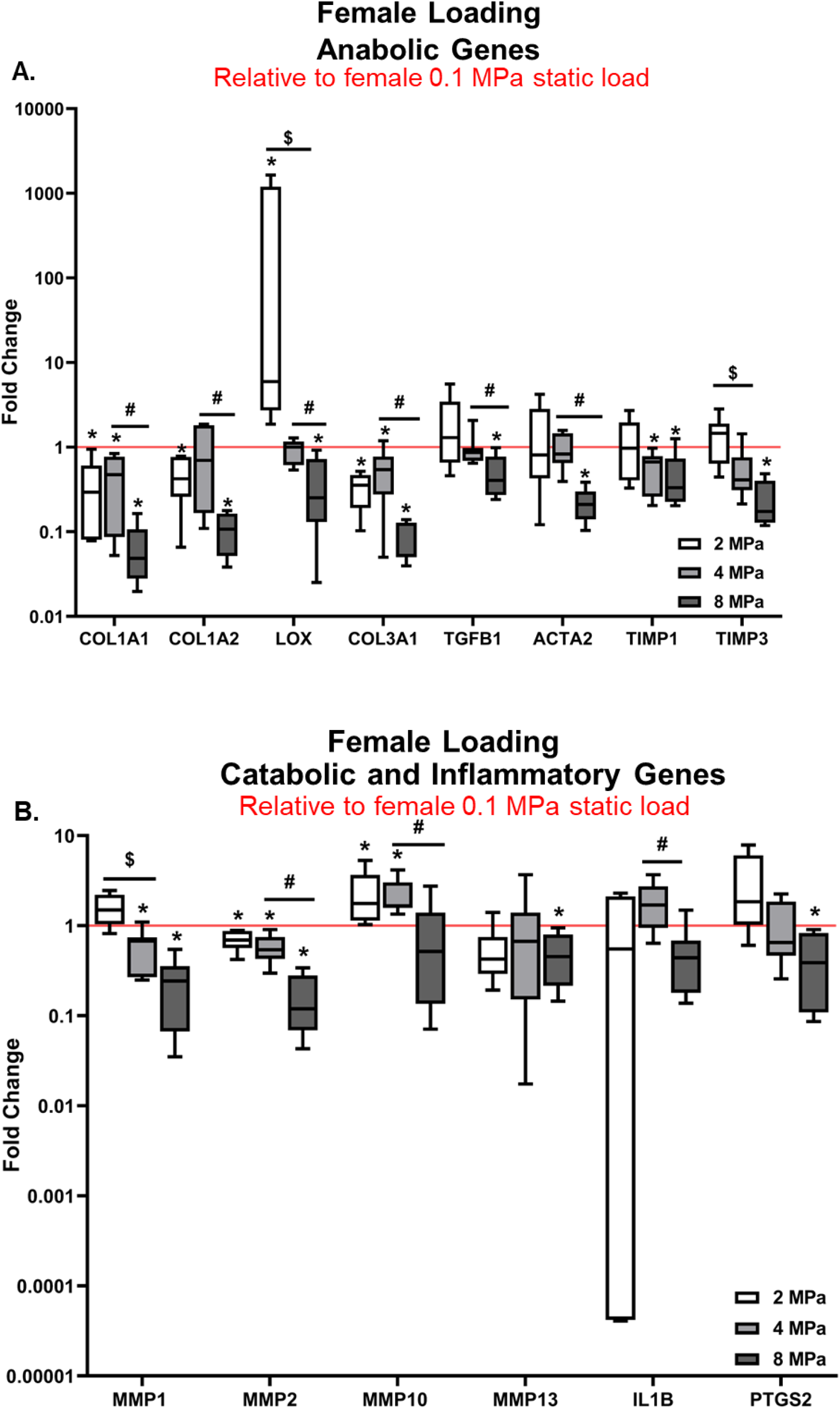
Female ACLs turn off gene expression with mechanical loading. RT-qPCR analysis of female ACLs after being cyclically loaded to 2, 4, and 8 MPa relative to the 0.1 MPa static load (represented by the red line at 1). (A) Anabolic markers. (B) Catabolic and inflammatory markers. (n = 6 for 2 MPa, n = 6-7 for 4 MPa, and n = 6-7 for 8 MPa) * p < 0.05 evaluated by one sample Wilcoxon test). $ represents statistical difference between 2 MPa and 4 MPa (p < 0.05 evaluated by Mann Whitney test). # represents statistical difference between 4 MPa and 8 MPa (p < 0.05 evaluated by Mann Whitney test). Data represented as box and whiskers plot with the whiskers representing the 10^th^ and 90^th^ percentile of data.

### Physical comparison of ACLs between sex

The average cross-sectional for female rabbit ACLs (3.1 ± 0.4 mm^2^) was significantly smaller than the male rabbit ACLs (3.5 ± 0.7 mm^2^) (p < 0.001) (**Fig. 5B**). Similarly, the length for female ACLs (7.5 mm ± 0.5 mm) was significantly shorter than the male ACLs (8.1 mm ± 0.9 mm) (p < 0.0001) (**Fig. 5C**). However, in all loading conditions, there was no significant difference in the macroscale strains between male and female ACLs (p = 0.73, p = 0.19, and p = 0.63) (**Fig. 5A**).

**Figure 5:**
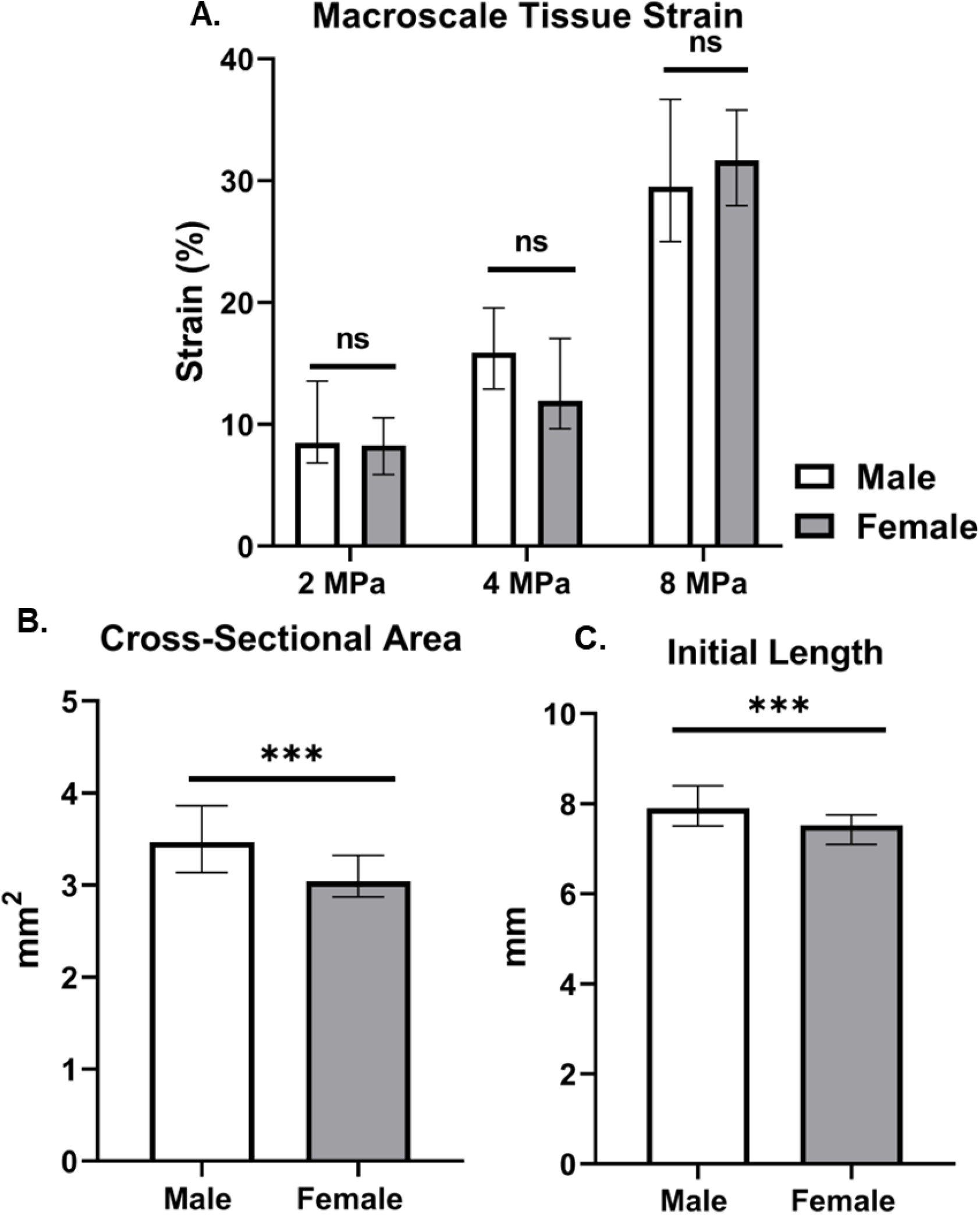
Female ACLs are smaller but exhibit similar tissue strains compared to male ACLs. (A) Macroscale tissue strain of male and female ACLs when cyclically loaded to 2, 4, and 8 MPa. (B) Cross-sectional area of male and female ACLs. (C) Initial length of male and female ACLs. (*** p < 0.001 evaluated by Mann-Whitney test). Macroscale data evaluated by a Mann Whitney test between sex at each loading condition. p > 0.05 at each load. Data represented as medians with error bars as the interquartile range.

## Discussion

The objective of this study was to investigate the remodeling response of ACLs to cyclic loading and to see if this response is sex specific. We hypothesized that ACLs would exhibit a dose response to cyclic loading, with lower load magnitudes initiating an anabolic response and higher loads initiating tissue degeneration. Consistent with our hypothesis, we found that tissue remodeling of the ACL was indeed dependent on the magnitude of cyclic loading. In the male ACL samples, upregulation of anabolic, catabolic, and inflammatory genes at 2 and 4 MPa suggests that tissue remodeling is activated with moderate cyclic loading. However, in contrast to our hypothesis, 8 MPa loading turned off tissue remodeling in male ACLs with downregulation of all gene categories. This suggests that the cellular response to cyclic mechanical loading is turned off at high physiological loads. Since these elevated loads produce microstructural damage within the ACL ^4,5^, a lack of cell-mediated tissue remodeling suggests that this damage is not repaired, thereby predisposing the ACL to injury. Our results are different from previous literature in tendon, which demonstrated that elevated cyclic loading initiates a catabolic cellular response ^30^. However, in both tendons and the ACL, elevated load is detrimental to tissue remodeling. Together, this supports the hypothesis that noncontact ACL injuries are attributed to fatigue-induced accumulation of microtrauma that predispose the ACL to injury, similar to the onset of tendon injuries ^31–34^

We also found that the remodeling response of ACL’s is sex specific. Male ACLs activate a tissue remodeling response at 2 and 4 MPa cyclic loads. In contrast, at all loading levels, female ACLs turn off tissue remodeling with downregulation of all gene categories. This was surprising given the fact that female ACLs exhibited a greater remodeling potential at baseline with significant upregulation of all gene categories compared to male ACLs. While downregulation of all gene categories was contrary to our hypothesis that female ACLs would exhibit an elevated catabolic response to high load magnitudes, our results still suggest that females have a reduced ability to repair tissue damage. While it is unclear whether the sex-specific effects we found in rabbits translates to humans, our results are in agreement with previous literature demonstrating that estrogen and mechanical loading inhibits the expression of collagen I in isolated porcine ACL fibroblasts ^35^. Given that this decreased remodeling potential is maintained across different species, it is possible that this could be maintained in humans as well. Together, our data suggest that the increased rate of ACL injuries in women is due to an impaired remodeling response to mechanical loading.

This study does have some limitations. First, while we selected our loading levels based on *in vivo* measurements of ACL loading ^36,37^, cyclic tensile loading doesn’t perfectly mimic the complicated loading environment of the ACL *in vivo.* Second, previous literature demonstrates that female human ACLs have inferior mechanical properties compared to male ACLs ^38^, which would create different cellular strains for the same applied stress. However, we found that there was no difference in the macroscale tissue strain between males and females at any load level, which suggests that the differences in gene expression observed between sexes were not due to differences in cellular strains. Third, 8 hours of loading is not enough time to induce fatigue damage or remodeling of the tissue structure, which is why we only measured changes in gene expression. Still, the observed changes in gene expression does provide insight into the cellular response to the different loading levels. Fourth, due to financial restrictions, our sample size was too small to correct for multiple comparisons in our statistical analysis. Nevertheless, the clear trends observed across loading levels and sample sex suggest that these data are accurate indicators of the biological response of ACLs to cyclic loading. Finally, it is important to note that rabbit ACLs have anatomical and biomechanical differences compared to human ACLs ^39^. Nevertheless, we found that female rabbit ACLs have a smaller cross-sectional area and length compared to male rabbit ACLs, which matches the size differences in male and female human data ^40,41^. Furthermore, the strain magnitudes created by physiological loading of our explants are consistent with the strain magnitudes observed in human ACLs ^37^.

In conclusion, this study demonstrates that the biological response of rabbit ACLs to cyclic loading is dependent on the load magnitude and is indeed sex specific. This study supports the novel hypothesis that noncontact ACL injury is a consequence of failed tissue remodeling and inadequate repair of microtrauma resulting from fatigue loading. Furthermore, our data demonstrate that female ACLs downregulate genes responsible for tissue remodeling in response to cyclic loading, which may help explain the increased rate of ACL tears in women. Future work will further investigate this hypothesis by measuring the structural and mechanical changes that occur in ACL explants loaded for longer durations. This will determine the effects of tissue remodeling in response to cyclic loading on tissue mechanics and directly compare the ability of male and female ACLs to repair fatigue damage. Additionally, our explant model enables us to identify the mechanisms underlying this sex-specific response to cyclic loading by perturbing mechanotransduction pathways via small molecule inhibitors and chemical agents. Together, these data can provide insight into the remodeling response of ACLs in vivo and potentially offer novel approaches for preventing ACL rupture.

## Acknowledgements

Funding for this project was provided by the Orthopeadic Research Foundation (234995) and the Congressionally Directed Research Program (W81XWH2110152). We also would like to acknowledge the Genomics core at Penn State for running the PCR plates. The authors have no conflict of interest.

