## Supplemental Figures 1-3 for "Cyclic Loading Induces Anabolic and Catabolic Gene Expression in ACLs in a Load-Dependent and Sex-Specific Manner"

**Figure S-1: Explant gene expression reaches a homeostasis after 18-hour overnight culture.** RT-qPCR analysis of male ACLs either cultured overnight or statically loaded to 0.1 MPa relative to freshly harvested ACLs (represented by the red line at 1). (A) Anabolic markers. (B) Catabolic and inflammatory markers. (n = 6 for overnight culture and n = 3-9 for 0.1 MPa static culture) * p < 0.05 evaluated by one sample Wilcoxon test. # represents statistical difference between overnight culture and 0.1 MPa static load (p < 0.05 evaluated by unpaired student t-test). Data represented as box and whiskers plot with the whiskers representing the 10^th^ and 90^th^ percentile of data.

**Figure S-2: Cell viability increases with tissue depth and is independent of sex.** (A) Cell viability as a function of tissue depth of male ACLs statically loaded to 0.1 MPa (n = 6). (B) Cell viability as a function of tissue depth of male ACLs cyclically loaded to 2 MPa (n = 7). (C) Cell viability as a function of tissue depth of male ACLs cyclically loaded to 4 MPa (n = 5). (D) Cell viability as a function of tissue depth of male ACLs cyclically loaded to 8 MPa (n = 5). (E) Comparison of male and female ACLs cyclically loaded to 4 and 8 MPa (n = 2 for female ACLs). Data is represented as median with interquartile range. Friedman test was conducted to determine significance across depth in each loading condition.

**Figure S-3: Male ACLs respond differently than female ACLs at each loading level.** RT-qPCR analysis of male and female ACLs freshly harvested after being cyclically loaded to 2, 4, and 8 MPa relative to their sex-matched 0.1 MPa static load (represented by the red line at 1). (A) Anabolic markers for ACLs cyclically loaded to 2 MPa. (B) Catabolic and inflammatory markers for ACLs cyclically loaded to 2 MPa. n = (6-9) for males and n = 6 for females. (C) Anabolic markers for ACLs cyclically loaded to 4 MPa. (D) Catabolic and inflammatory markers for ACLs cyclically loaded to 4 MPa. n = (5-6) for males and n = (6-7) for females. (E) Anabolic markers for ACLs cyclically loaded to 8 MPa. (F) Catabolic and inflammatory markers for ACLs cyclically loaded to 8 MPa. n = 6 for males and n = (6-7) for females.* represents statistical differential expression (p < 0.05 evaluated by one sample Wilcoxon test). # represents statistical difference between 2 male and female ACLs (p < 0.05 evaluated Mann Whitney test). Data represented as box and whiskers plot with the whiskers representing the 10^th^ and 90^th^ percentile of data.

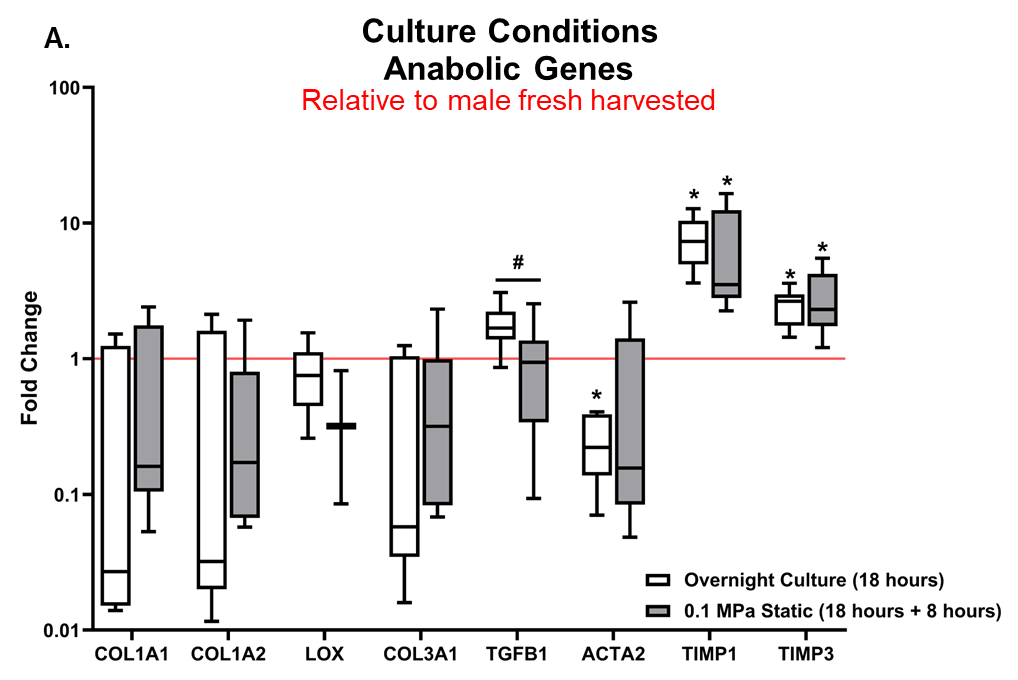

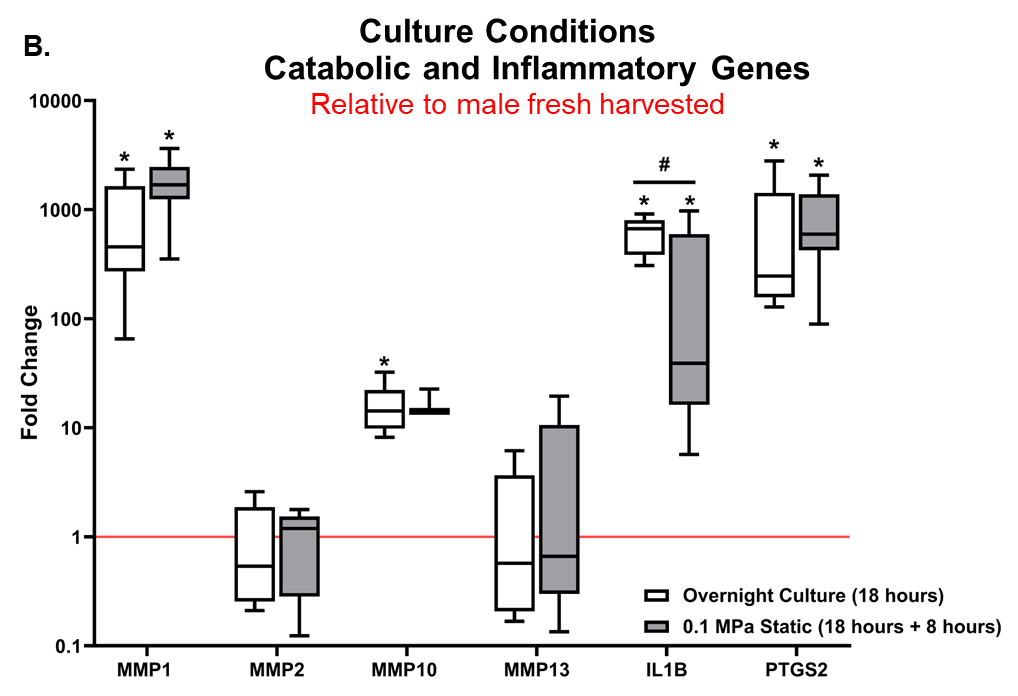

**Figure S-1**

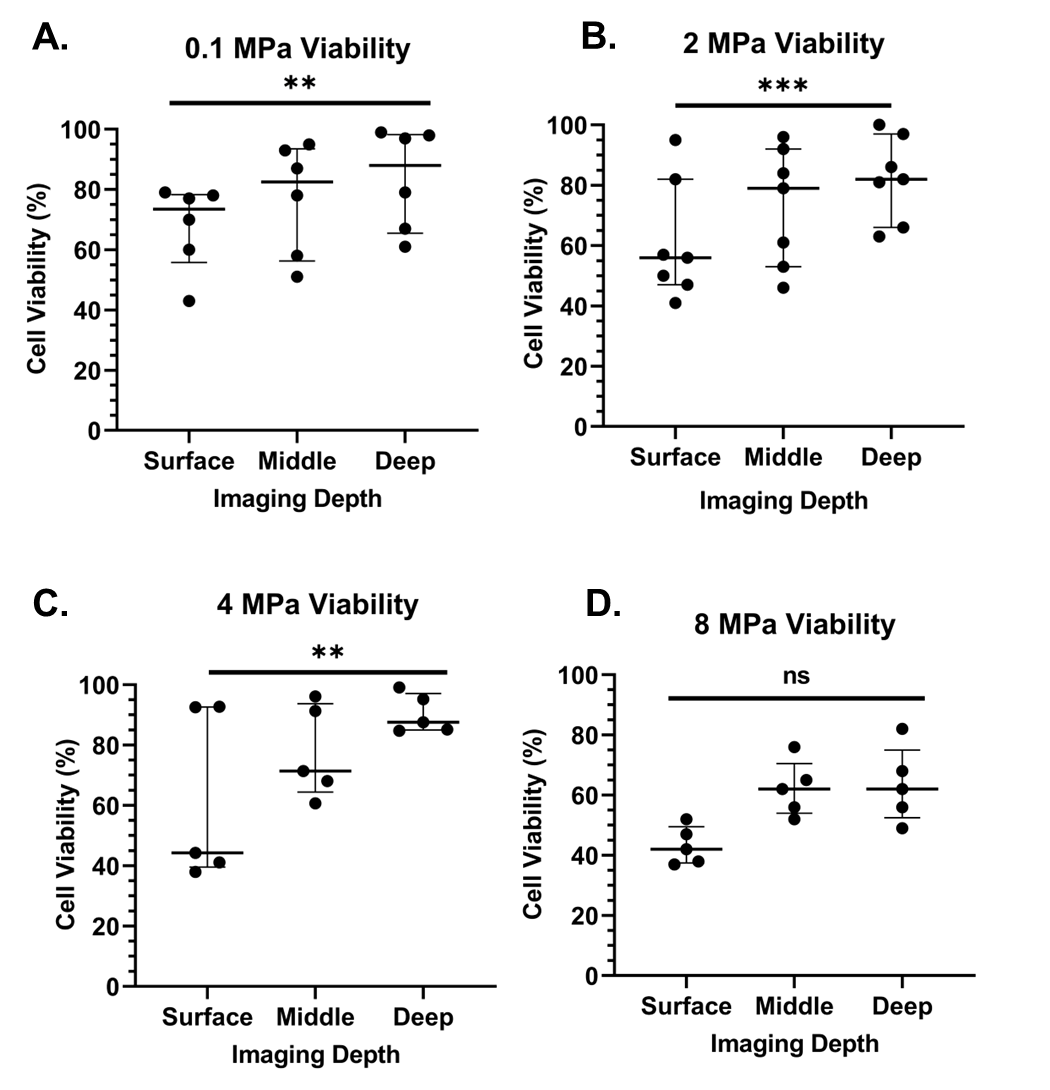

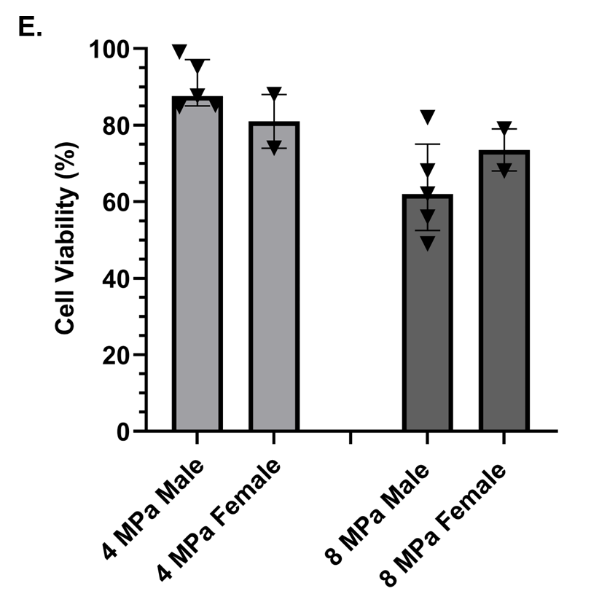

**Figure S-2**

**Figure S-3**

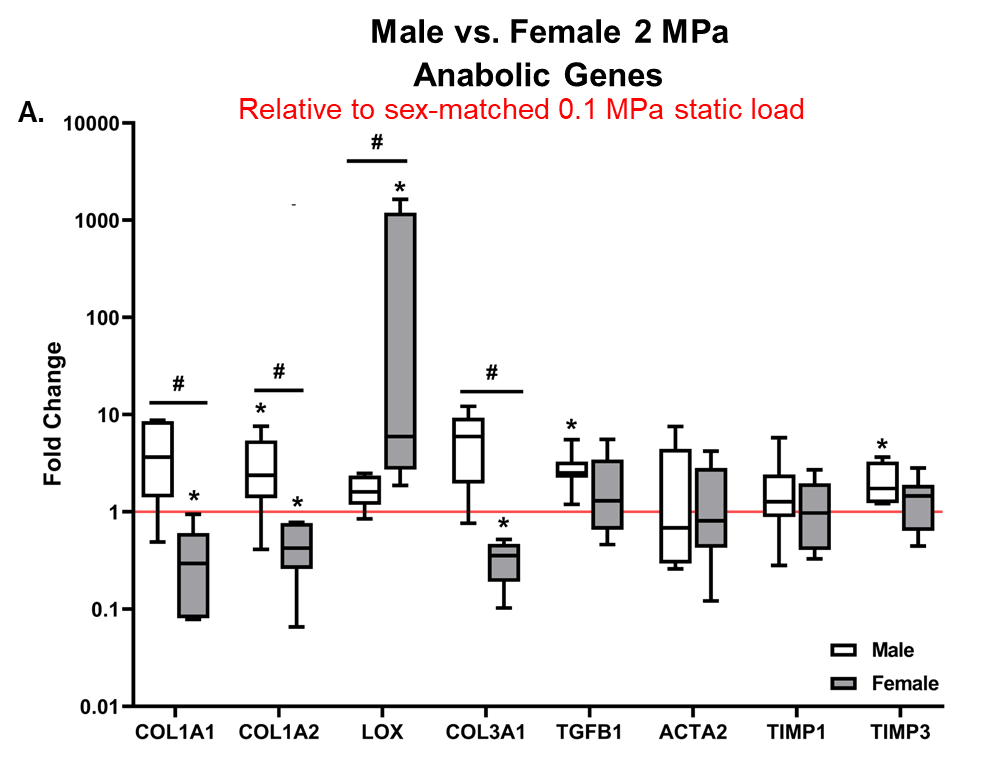

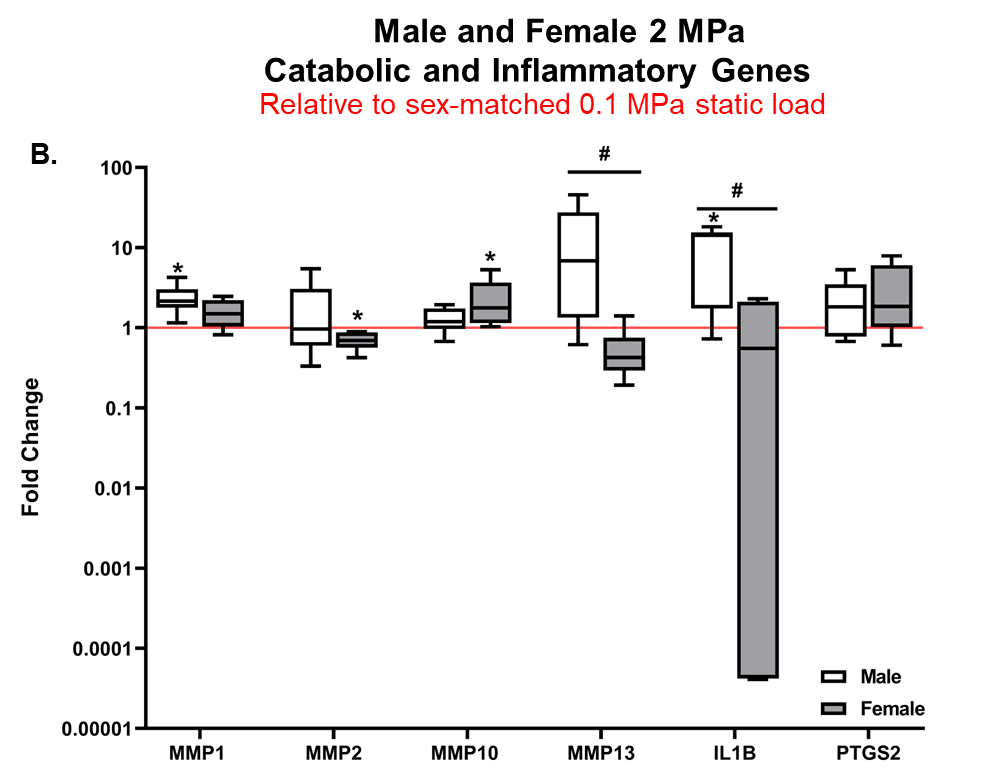

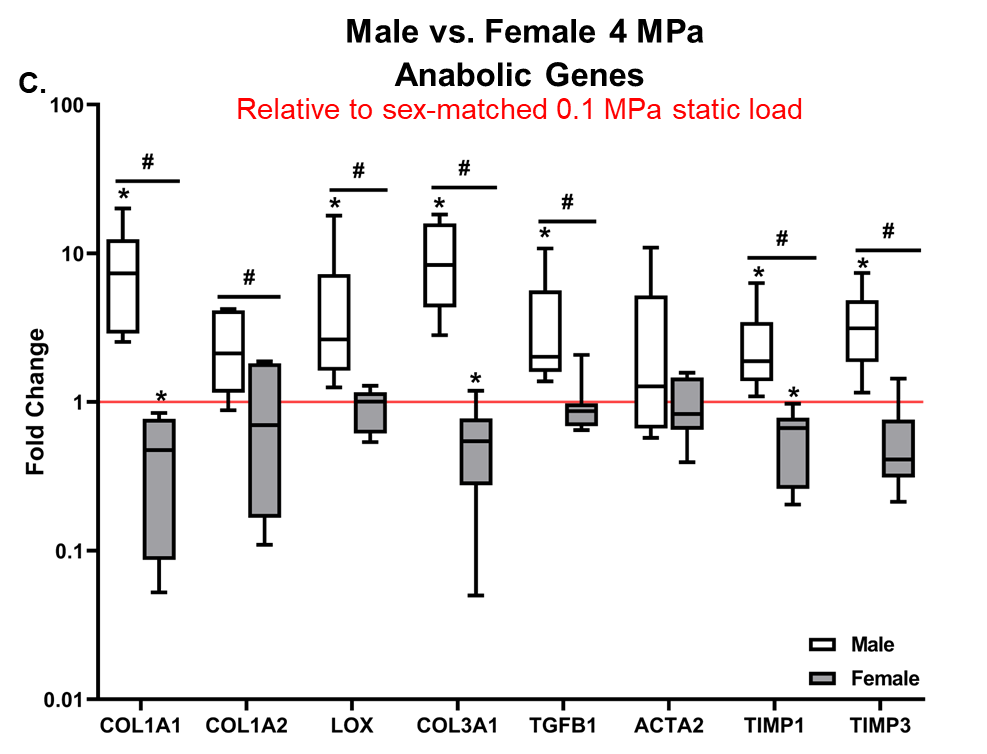

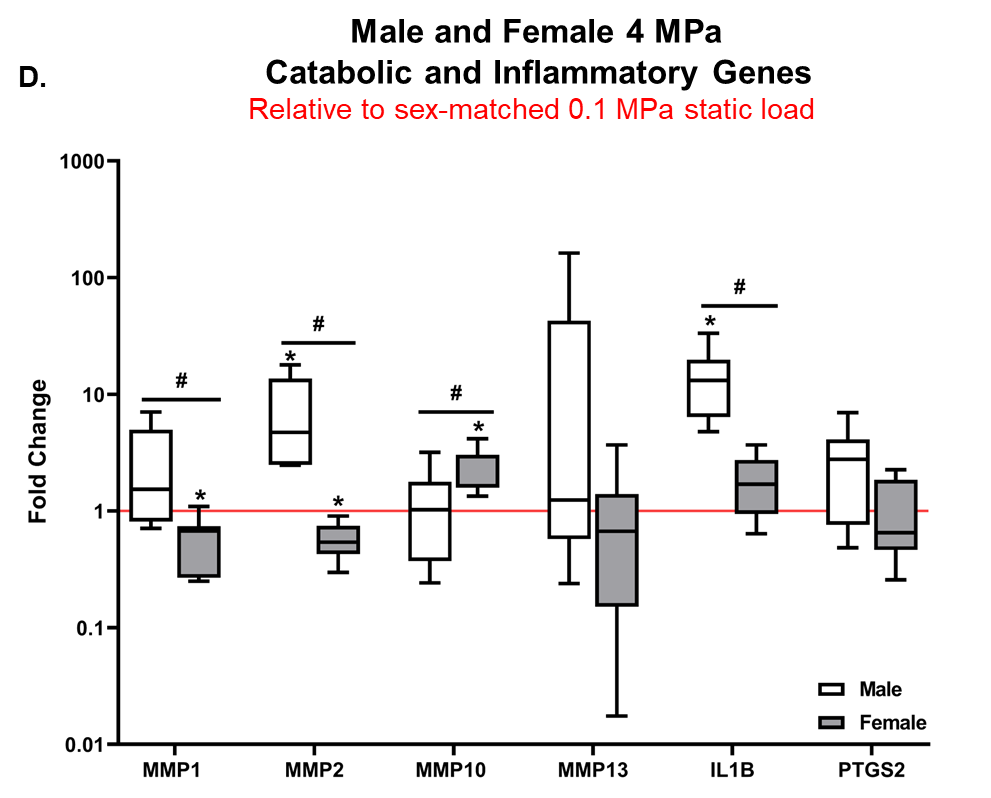

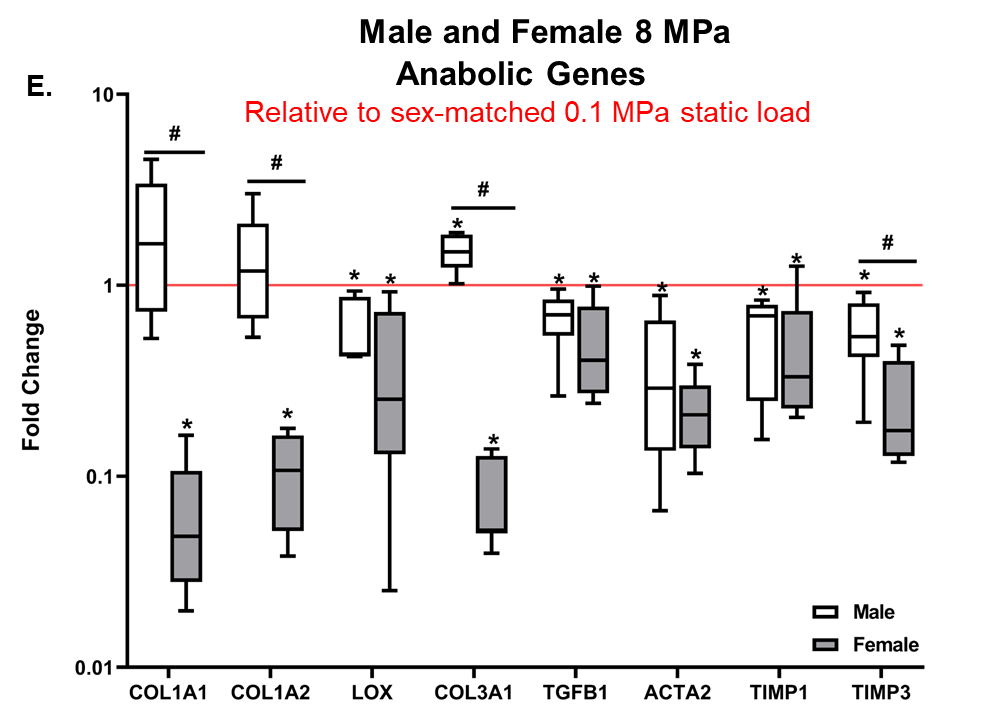

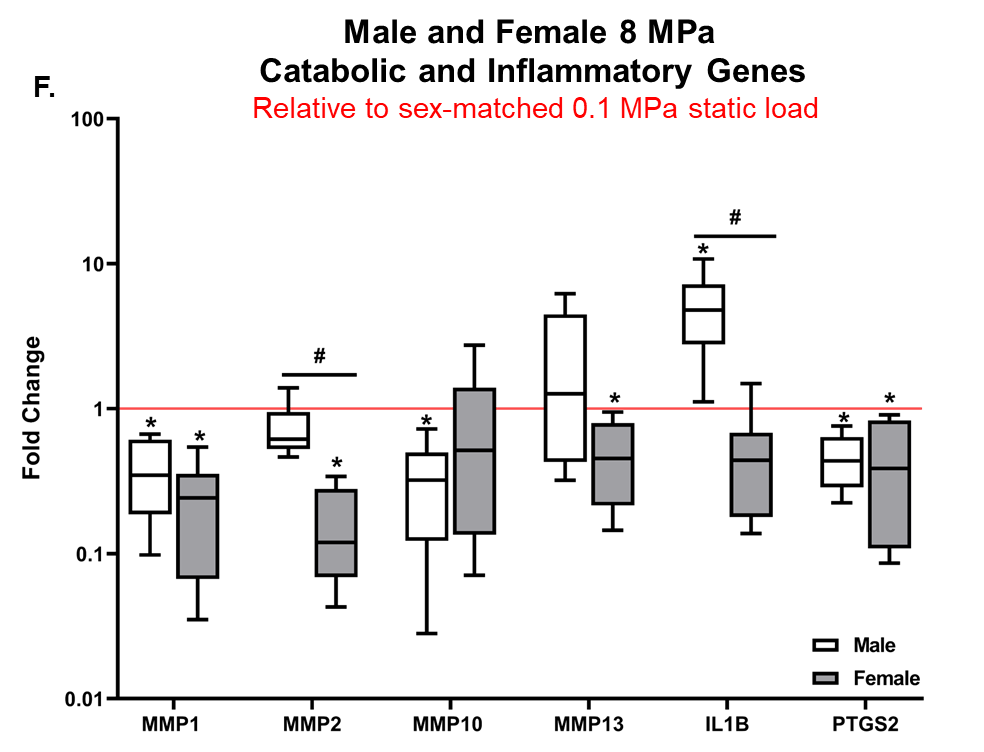
